# NF-YA Transcriptionally Activates the Expression of SOX2 in Cervical Cancer Stem Cells

**DOI:** 10.1101/599613

**Authors:** Wen-Ting Yang, Zong-Xia Zhao, Bin Li, Peng-Sheng Zheng

## Abstract

Roles for SOX2 have been extensively studied in several types of cancer, including colorectal cancer, glioblastoma and breast cancer, with particular emphasis placed on the roles of SOX2 in cancer stem cell. Our previous study identified SOX2 as a marker in cervical cancer stem cells driven by a full promoter element of SOX2 EGFP reporter. Here, dual-luciferase reporter and mutagenesis analyses were employed, identifying key cis-elements in the SOX2 promoter, including binding sites for SOX2, OCT4 and NF-YA factors in SOX2 promoter. Mutagenesis analysis provided additional evidence to show that one high affinity-binding domain CCAAT box was precisely recognized and bound by the transcription factor NF-YA. Furthermore, overexpression of NF-YA in primitive cervical cancer cells SiHa and C33A significantly activated the transcription and the protein expression of SOX2. Collectively, our data identified NF-YA box CCAAT as a key cis-element in the SOX2 promoter, suggesting that NF-YA is a potent cellular regulator in the maintenance of SOX2-positive cervical cancer stem cell by specific transcriptional activation of SOX2.

## Introduction

Tumor growth, metastasis and recurrence are driven by a small sub-population of cancer stem cells (CSCs)[1]. Most CSCs assays have thus far depended on a variety of different cell surface markers, including CD133, CD44, CD166, CD24 and so on[2]. However, surface markers can only be used to isolate the most common CSCs and these markers are often unstable in many somatic cancers[3]. Additionally, the results obtained with CSCs isolated using the same surface marker are not consistent among laboratories. Due to the instability and scarcity of surface markers in solid tumors, other methodological strategies have been widely explored to identify and isolate CSCs, including nuclear markers[4], side population phenotypes[5], sphere formation, and aldehyde dehydrogenase (ALDH) activity assays[6].

In our previous study, we have identified the expression of the embryonic stem cell-specific transcription factor SOX2 in primary cervical cancer tissues and tumorspheres formed by primary cervical carcinoma cells and found that SOX2 functions as an oncogene in cervical carcinogenesis by promoting cell growth and tumorigenicity, which was consistent with other lab’s results[7, 8]. Additionally, SOX2 is a key factor that controls the pluripotency, self-renewal and proliferation of embryonic stem cells[9]. It has been shown that murine and human embryonic and neural stem cells have high activity of SOX2[10, 11]. An increase in the expression of SOX2 has also been found in breast and glioblastoma CSC populations[12–14]. Based on these findings, cervical CSCs have been isolated and identified by sorting the endogenous SOX2-positive cells using a plasmid pSOX2/EGFP that contained the full length of SOX2 promoter with 11.5kb nucleotides positioned upstream of the EGFP reporter[7]. However, This plasmid was so large (approximately 16kb) that the transfection efficiency in cervical cancer cell lines was so low for sorting the endogenous SOX2-positive cells. Here, in order to explore the functional domain in the promoter of SOX2 helps us to efficiently capture CSCs and elucidate the role of SOX2, upstream regulatory factor, and related signaling pathways, we try to obtained the trans-factors that specifically bond to the key domain region of SOX2 promoter and activated the expression of SOX2

## Materials and methods

### Ethics Statement

Investigation has been conducted in accordance with the ethical standards and according to the Declaration of Helsinki and according to national and international guidelines and has been approved by the review board of the First Affiliated Hospital of Xi’an Jiaotong University.

### Cell lines and culture conditions

The human cervical cancer cell lines SiHa and C33A were obtained from the American Type Culture Collection (ATCC; Manassas, VA). SiHa and C33A cells were cultured in Dulbecco’s Modified Eagle Medium-high glucose (DMEM; Sigma-Aldrich, St. Louis, MO) supplemented with 10% fetal bovine serum (FBS; Invitrogen, Carlsbad, CA) and maintained at 37°C in an atmosphere containing 5% carbon dioxide.

### Flow Cytometry and Separation of cervical cancer stem cell by FACS

To obtain the EGFP^+^ and EGFP^−^ populations, SiHa and C33A cells were transfected with pSox2/EGFP plasmid using Lipofectamine 2000 (Invitrogen). Selection was performed using standard culture medium with 1mg/mL G418. The generation of single cell-derived cultures was performed using a FACSAria (Becton Dickinson, Franklin Lakes, NJ). The sorting gates were established as the highest and lowest 10% of the EGFP-expressing cells. The cells were cultured in DMEM/F12 with N2 and B27 supplements (Invitrogen), 20ng/mL human recombinant epidermal growth factor (EGF) and 20ng/mL basic broblastic growth factor (bFGF; PeproTech Inc., Rocky Hill, NJ).

### Construction of SOX2 promoter luciferase reporter

Luciferase reporter plasmids with a pGL3 backbone (Promega Corporation, catalog number: E1751) have been utilized to characterize the transcriptional effects of mutations in the SOX2 promoter. We redesigned the pGL3 basic vector by cloning the 5’ UTR (110bp) and 3’ UTR (1264) regions of the SOX2 promoter though 5’ and 3’ UTRs on both sides of the luciferase gene using an In-Fusion PCR Cloning Kit. The resulting vector was designated PGL3-pSox2 mini (Takara Bio Inc, Dalian, China).

Next, the deletion plasmids including PGL3-pSox2-4650+1828, PGL3-pSox2-3081+1828, PGL3-pSox2-1185+1828, PGL3-pSox2 mini+1828, PGL3-pSox2-4650, PGL3-pSox2-3081, PGL3-pSox2-1185, PGL3-pSox2-1185+1600, PGL3-pSox2-1185+1200 and PGL3-pSox2-1185+600 for the SOX2 promoter were constructed from the phSox2/EGFP vector that was used to sort the SOX2-positive cervical CSCs in our previous study. The predictive binding domains in SOX2 promoter were mutated by PCR. The primers used for constructing the deletions and mutations in the SOX2 promoter are listed in STable 1

### Dual luciferase reporter assay

SOX2 promoter luciferase reporters and pTK-RL plasmids were transiently co-transfected into tumor cells (5×10^4^) plated in the 24-well plate dish, while the activity of both firefly and Renilla luciferase reporters was determined 48 hours post transfection using the Dual Luciferase Assay kit (Promega, Madison, WI, USA), according to the manufacturer’s instructions. The SOX2 promoter luciferase reporter activity was presented as the relative ratio of firefly luciferase activity to Renilla luciferase activity. The specific activity was dis-played as the fold change of the experimental group versus the control group. All experiments were performed in triplicate.

### Western blotting

Western blot analyses were performed as previously described using 30ug cell lysates. The primary antibodies were goat polyclonal anti-SOX2 (1:500, Santa Cruz, CA, USA), rabbit polyclonal anti-OCT4 (1:1000, Santa Cruz, CA, USA), mouse anti-NF-YA (1:1000, Santa Cruz, CA, USA) and GAPDH (1:1000, Santa Cruz, CA, USA), The secondary incubation antibodies used a horseradish peroxidase-conjugated anti-rabbit, anti-mouse or anti-goat IgG (Thermo Fisher Scientific, New York, NY, USA). The signals were then detected by enhanced chemiluminescence reagent (Millipore, Billerica, MA, USA).

### Immunofluorescence

Cells were cultured on glass coverslips for 48 hours, fixed in 4% paraformaldehyde for 30 minutes at room temperature, and thenpermeabilized with 0.1% Triton X-100 for 20 minutes at room temperature (Sigma, St. Louis, MO). Anti-rabbit Alexa Fluor 488 and antigoat Alexa Fluor 555 were purchased from Invitrogen, and 4’, 6-diamidino-2-phenylindole (DAPI) was bought from Sigma. Co-localization of SOX2 and OCT4 was analyzed using a Leica TCS SP5 confocal microscope (Leica TCS SP5, Wetzlar, German). Images were captured with a Leica DFC 500 digital camera and processed with LAS AF software (Leica).

### Statistical analysis

Statistical analyses were performed using GraphPad Prism 5.01 software (La Jolla, CA, USA). In comparisons of 2 groups, Two-taild student’s *t*-test was used to determine the statistical significance. To examine differences among 3 groups, an ANOVA was performed. Kaplan–Meier survival analysis was performed and survival curve comparisons were performed using the log-rank (Mantel-Cox) test. A *p* value of < 0.05 was regarded as statistically significant.

## Results

### An integrated analysis of SOX2 transcription element in cervical CSCs

In our previous study[10], we constructed the plasmid pSOX2/EGFP, which contains 11.5 kb of the human SOX2 promoter sequence positioned upstream of the enhanced green fluorescent protein (EGFP) reporter. This plasmid also contains the human Sox2 3’UTR, 3’poly (A) tail, and 3’enhancer positioned downstream of the EGFP reporter. Then, SOX2-positive cervical cancer cells were isolated by pSOX2/EGFP plasmid in cervical cancer cell lines, such as SiHa and C33A, and confirmed that SOX2-positive subpopulation cells exhibited dominant characteristics of CSCs, including tumorigenicity, self-renewal and differentiation. Here, the key cis-element region in the SOX2 promoter and the upstream trans-factors that activated the expression of SOX2 were both explored.

Firstly, in order to obtain the dominant region driving the expression of SOX2, we analyzed the full length of SOX2 promoter (Fig. 1a), which contained 4650bp and 1828bp though two sides of SOX2 CDS region. Then, several deletions were fused to pGL3 basic luciferase reporter plasmids through Web Promoter Scan Service and TFSEARCH on line software (fig. 1b). Then, the optimized response element was determined by dual-luciferase reporter assay in SOX2-positive and -negative cervical cells from SiHa, and we found that the luciferase activity of reporters containing −1185bp region in SOX2-EGFP+ cells was significantly higher than that in SOX2-EGFP-cells regardless of whether there was −1828 downstream or not (Fig.1c, *p* < 0.05). Furthermore, we found that the activity of pSOX2-1185-luc-1828 was much higher than that of pSOX2-1185-luc (*p* < 0.05). We supposed that the +1828bp in the 3’ UTR region containing two binding sites of SOX2 and OCT4 might function in the transcription activity of SOX2 promoter. According to constructing the mutations, deleting the two binding sites significantly decreased the luciferase activity comparing with pSOX2-1185-luc-1828 reporter. However, the individual 3’ UTR region including SOX2 and OCT4 binding sites (pSOX2-mini-luc-1828) could not activate the transcription at all. Thus, we hypothesized that the −1185bp to +1828bp region (pSOX2-1185-luc-1828) of the SOX2 promoter may be necessary for transcriptional activation of SOX2 in cervical CSCs.

**Figure 1.**
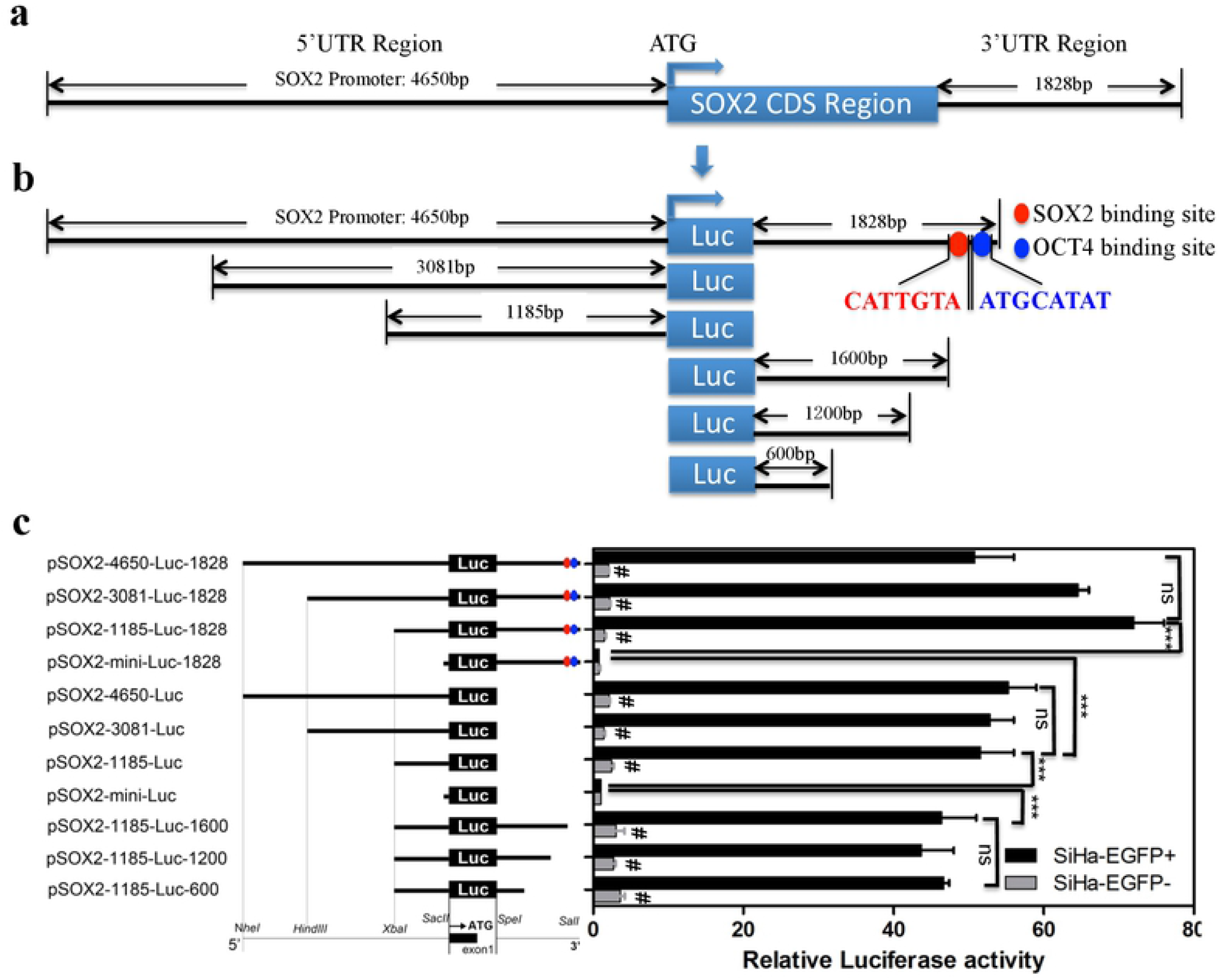
An integrated analysis of the SOX2 transcriptional element in cervical CSCs. (a and b) The diagram of full promoter of SOX2 and the deletions containing the possible cis-acting elements and the SOX2 and OCT4 binding sites (c) The full promoter of SOX2 (pSOX2-4650-luc-1828) and deletions were constructed and luciferase activity relative to Renilla control was measured in SiHa-EGFP+ and SiHa-EGFP-cells. The transcriptional activity of *pSOX2-mini-luc* served as the negative control and the SOX2 transcriptional activity was expressed relative to pSOX2-mini-luc. Data is presented as the mean ± SD of experiments in triplicate and statistically analyzed with student’s *t*-test. The symbol *** represents a *p* value of < 0.001 while ns indicates no statistical difference.

### OCT4 partially increased the transcriptional activation of the SOX2 promoter

In order to confirm that the transcription factor SOX2 and OCT4 were necessary for the expression of SOX2, which has been confirmed in the previously report, we deleted or mutated the two binding sites in the 3’UTR region of SOX2 promoter and changed the location of the sites within the upstream region of the SOX2 promoter. The mutation sequences (SOX2: from CATTGTA mutated to ACGGTGC and OCT4: from ATGCATAT mutated to CGTACGCG) of SOX2 (Red) and OCT4 (Blue) binding sites are shown in Fig. 2a. The results from the dual-luciferase assay showed that either or both mutations of SOX2 and OCT4 binding sites significantly inhibited the luciferase activity (*p* < 0.05, Figure 2b) in SOX2-positive cervical CSCs. Moreover, the two binding site regions of SOX2 (S) and OCT4 (O) (SO region: CATTGTA and ATGCATAT) joined directly to the 3’ UTR terminus in the plasmid pSOX2-1185-luc showed equal transcriptional activity to the pSOX2-1185-luc-1828 plasmid. However, inserting the SO sequence into 5’UTR region (pSOX2-SO-1185-luc and pSOX2-1185SO-luc), the luciferase activity in SOX2-positive+ cells not only did it increased, but it was restrained compared with pSOX2-1185-luc-SO reporter (*p* < 0.05)

**Figure 2.**
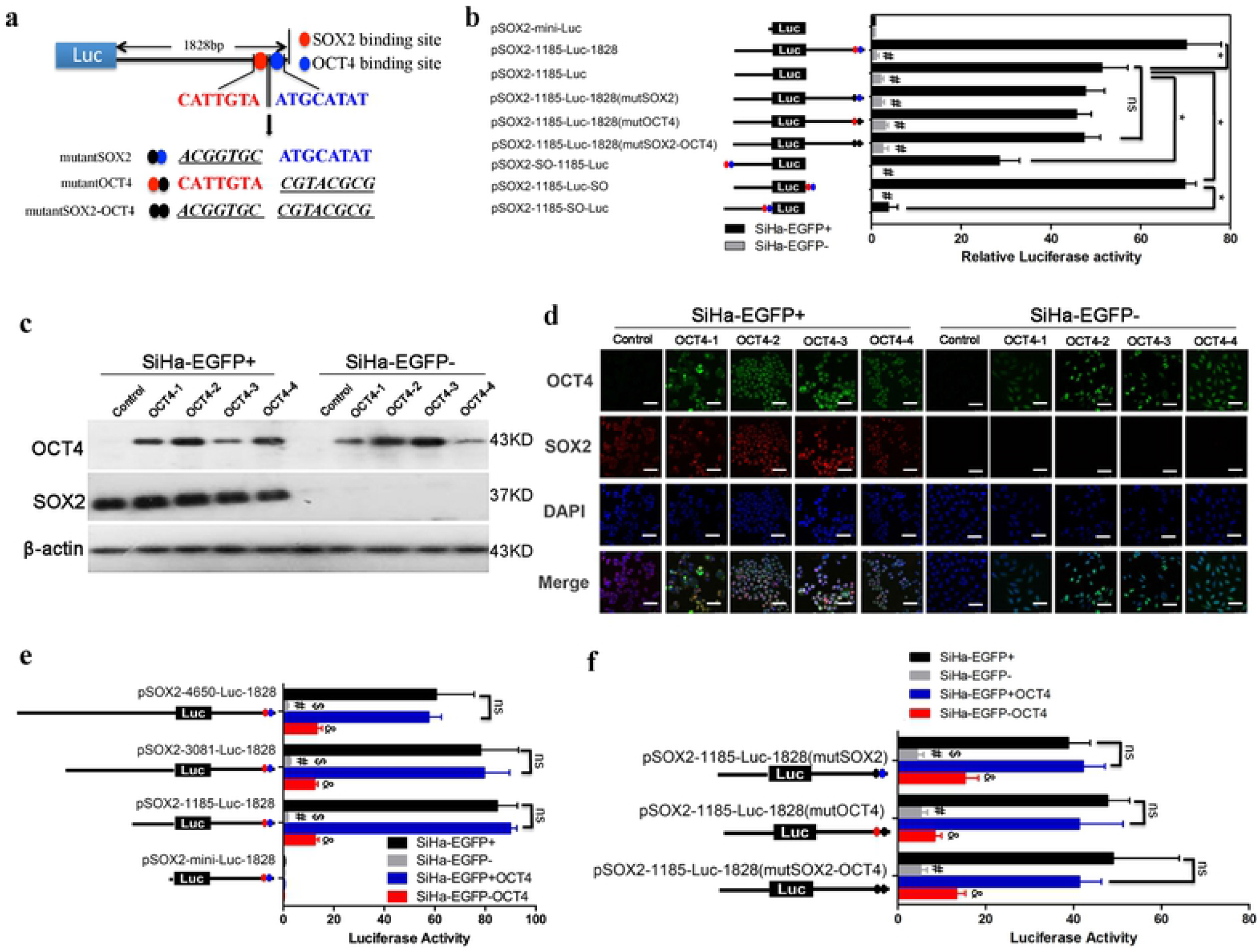
OCT4 partial increased SOX2 transcriptional activation. (a) The mutations of SOX2 and OCT4 binding sites downstream of the SOX2 promoter 3’UTR. (b) The luciferase activity of mutations in SOX2 promoter was detected. (S: SOX2; O: OCT4; red dot: SOX2 binding site; blue dot: OCT4 binding site; black dot: mutant of SOX2 or OCT4 binding site). The expression level of OCT4 was detected in OCT4-overexpressing SiHa-EGFP+ and SiHa-EGFP-cells by Western blot (c) and immunofluorescence (d). (e and f) The luciferase activity of the SOX2 promoter deletions and mutations in OCT4-overexpressing cells. Data is presented as the mean ± SD of experiments in triplicate and statistically analyzed using student’s t-test. The symbols represent the following: *, *p* < 0.05; #, SiHa-EGFP+ *vs* SiHa-EGFP- and *p* < 0.05; $, SiHa-EGFP- *vs* SiHa-EGFP-OCT4 and *p* < 0.05; &, SiHa-EGFP+OCT4 vs SiHa-EGFP-OCT4 and *p* < 0.05; ns = no statistical difference.

Next, the OCT4 protein was exogenously expressed in SOX2-positive and SOX2-negative SiHa cells. The expression level was detected by Western blot and immunofluorescence (Fig. 2c and d). Interestingly, the overexpression of OCT4 in SOX2-negative cells did not induce the expression of the SOX2 protein, which suggested that OCT4 could increase the transcription of SOX2 rather than trigger.

Furthermore, the transcription activation levels of OX2 were monitored by dual-luciferase assay in OCT4-overexpressing cells. OCT4 slightly increased the activation of transcription in all plasmids containing ~+1185 and ~-1828 regions, but without any effect in the plasmid only containing ~-1828 region (Fig. 2e). Additionally, OCT4 promoted the transcription of SOX2 in SOX2-negative cells with the plasmid pSOX2-1185-Luc-1828mutSOX2 that had a mutation in the SOX2 binding site. However, when the OCT4 binding site was mutated as with plasmid pSOX2-1185-Luc-1828mutOCT4, the transcription of SOX2 was not significantly altered (Fig. 2f). These results suggest that the OCT4 binding site in the SOX2 promoter could improve the transcriptional activation but was not required for SOX2 expression.

### The NF-YA binding site CCAAT/ATTGG box was required for the transcription of SOX2 in cervical CSCs

All these results above suppressed that the upstream ~+1185 region of SOX2 promoter could be of crucial importance for the transcription of SOX2. According to the functional analysis of transcription factor binding sites, we found two binding sites of SOX2 and NF-YA in the candidate region. Then, mutations in plasmid pSOX2-1185-luc were constructed to harbor mutations from GAACAATA to TCCACCGT in the SOX2 binding site and from TGATTGGTC to GTCGGTTGA in the NF-YA binding site (Fig. 3a). The luciferase activity of either SOX2 or NF-YA mutation in SOX2-positive cells was significantly inhibited compared with pSOX2-1185-Luc reporter (Fig. 3b). Of note the mutation in the NF-YA binding site in SOX2-positive cells resulted in a decrease of transcriptional activity to the level of that in SOX2-negative cells (Fig. 3b). This suggests that CCAAT/ATTGG box located −485 upstream of SOX2 promoter was indispensable for the transcription of SOX2.

**Figure 3.**
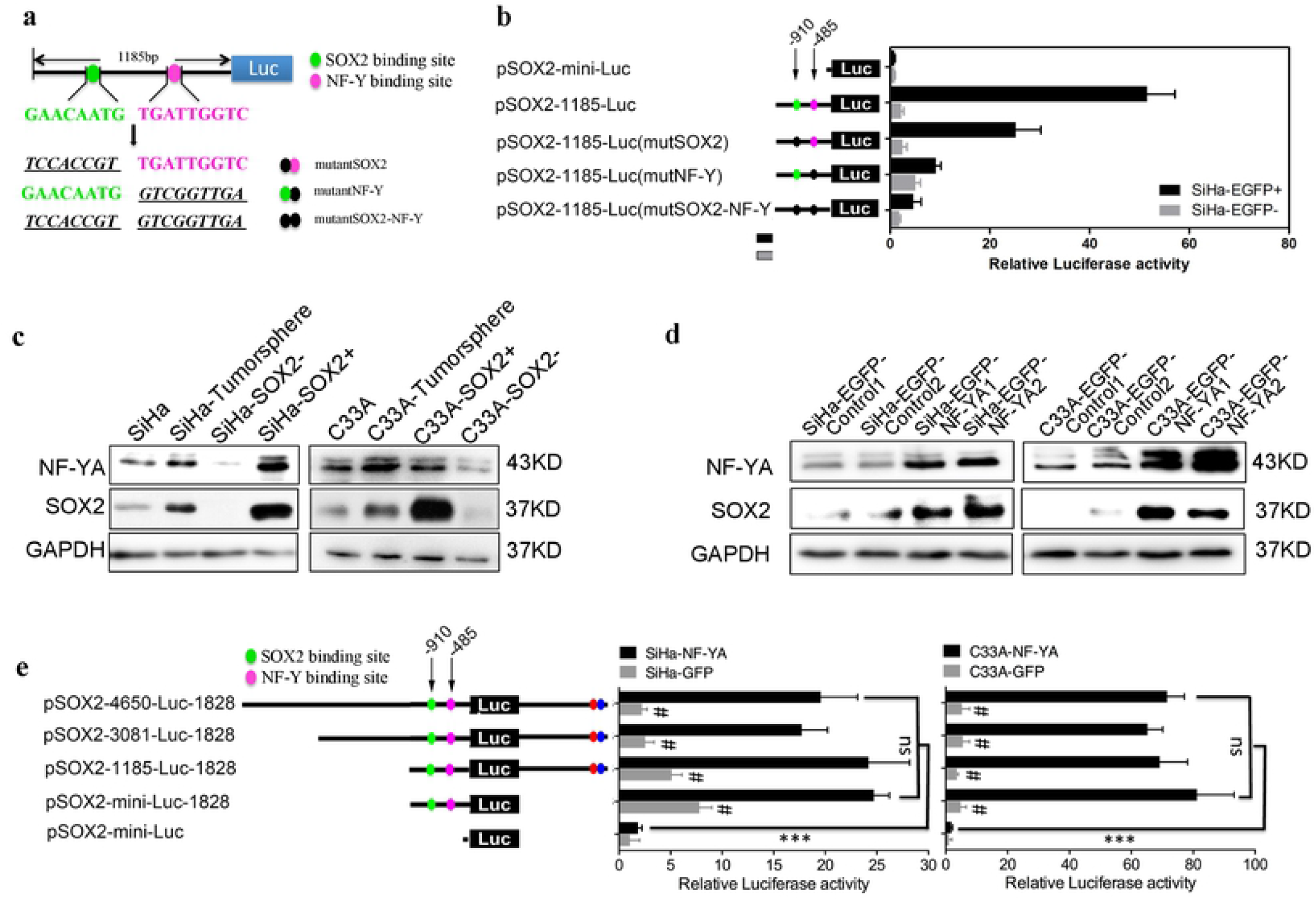
The NF-YA binding site CCAAT/ATTGG box was required for the transcription of SOX2 in cervical CSCs. (a) The mutations of SOX2 and NF-Y binding sites upstream of the SOX2 promoter 5’UTR. (b) The luciferase activity of the SOX2 promoter mutations (green dot: SOX2 binding site; pink dot: NF-Y binding site; black dot: mutant of SOX2 or NF-Y binding site). (c and d) NF-YA protein expression was detected in SiHa and C33A cells cultured in adherent culture and tumorsphere, as well as SOX2+ and SOX-SiHa and C33A cells. NF-YA was overexpressed in SiHa and C33A cells and the expression levels of NF-YA, SOX2 and OCT4 were detected by Western blot. (e) The luciferase activity of the SOX2 promoter deletions in NF-YA-overexpressing SiHa and C333A cells. Data is presented as the mean ± SD of experiments in triplicate and statistically analyzed using student’s *t*-test. The symbols represent the following: ***, *p* < 0.001; #, NF-YA group *vs* GFP group and *p* < 0.05; ns = no statistical difference.

The CCAAT/ATTGG box was specifically bound by transcript factor NF-YA, suggesting that NF-YA might play an important role in the maintenance of cervical CSCs properties derived from SOX2 transcriptional activity. Thus, we detected the expression of NF-YA in isolated SOX2-positive and -negative SiHa and C33A cells. The expression level of NF-YA protein in SOX2-positive cells was higher that that in SOX2-negative cells both in SiHa and C33A cells (Fig. 3c). Fortunately, tumorspheres cultured in serum-free media also showed both higher SOX2 and NF-YA expression than they adherent cultured (Fig. 3c). In order to confirm that the CCAAT/ATTGG box was vital to the transcriptional activation of SOX2 by the factor NF-YA, we exogenously overexpressed NF-YA in SOX2-negative SiHa and C33A cells and found by western blot that SOX2 was upregulated (Fig. 3d). OCT4 as the downstream gene of SOX2 was also upregulated in NF-YA overexpressed SOX2-negative cells compared with that in SOX2-negative cells, which might be induced by the upregulation of SOX2. Meanwhile, the transcript level of SOX2 driven by the plasmids containing the ~+1185 region was also significantly improved in NF-YA-overexpressed cells (Fig. 3e, *p* < 0.05). These results suggest that the CCAAT/ATTGG box in the SOX2 promoter region was necessary for transcriptional activation of SOX2 with its trans-factor NF-YA upregulating the expression of the SOX2 protein.

### NF-Y specifically binds to CCAAT/ATTGG box upstream of SOX2 promoter in cervical CSCs

Since NF-YA, as a universally accepted trans-acting factor of the CCAAT/ATTGG box, could increase the expression of SOX2 in cervical CSCs, we investigated whether NF-YA could transcriptionally activate the expression of SOX2 via physiological binding to the cis-element of CCAAT/ATTGG box in SOX2 promoter. Here, NF-YA overexpressed SOX2-negative SiHa and C33A cells showed increased luciferase activity of pSOX-1185-luc as compared with SOX2-negative cells, respectively. However, when mutations were introduced to the CCAAT/ATTGG box site, there was no significant difference in luciferase activity between NF-YA-overexpressing SOX2-negative cells and SOX2-negative cells (Fig. 4a, *p* < 0.05). Moreover, in the deletions of the ~1185 region, only pSOX2-SN1-Luc, including both SOX2 and NF-YA binding sites and the unknown middle region, could be transcriptionally activated in NF-YA-overexpressing SOX2-negative SiHa and C33A cells (Fig. 4b, *p* < 0.05). Additionally, SOX2 was silenced in NF-YA over expressing SOX2-negative C33A cells and the luciferase activity was significantly reduced compared with NF-YA overexpressing cells (Fig. 4c and d). The findings suggest that that NF-Y transcriptionally activated the expression of SOX2 in cervical CSCs by binding to the CCAAT/ATTGG box upstream of the SOX2 promoter.

**Figure 4.**
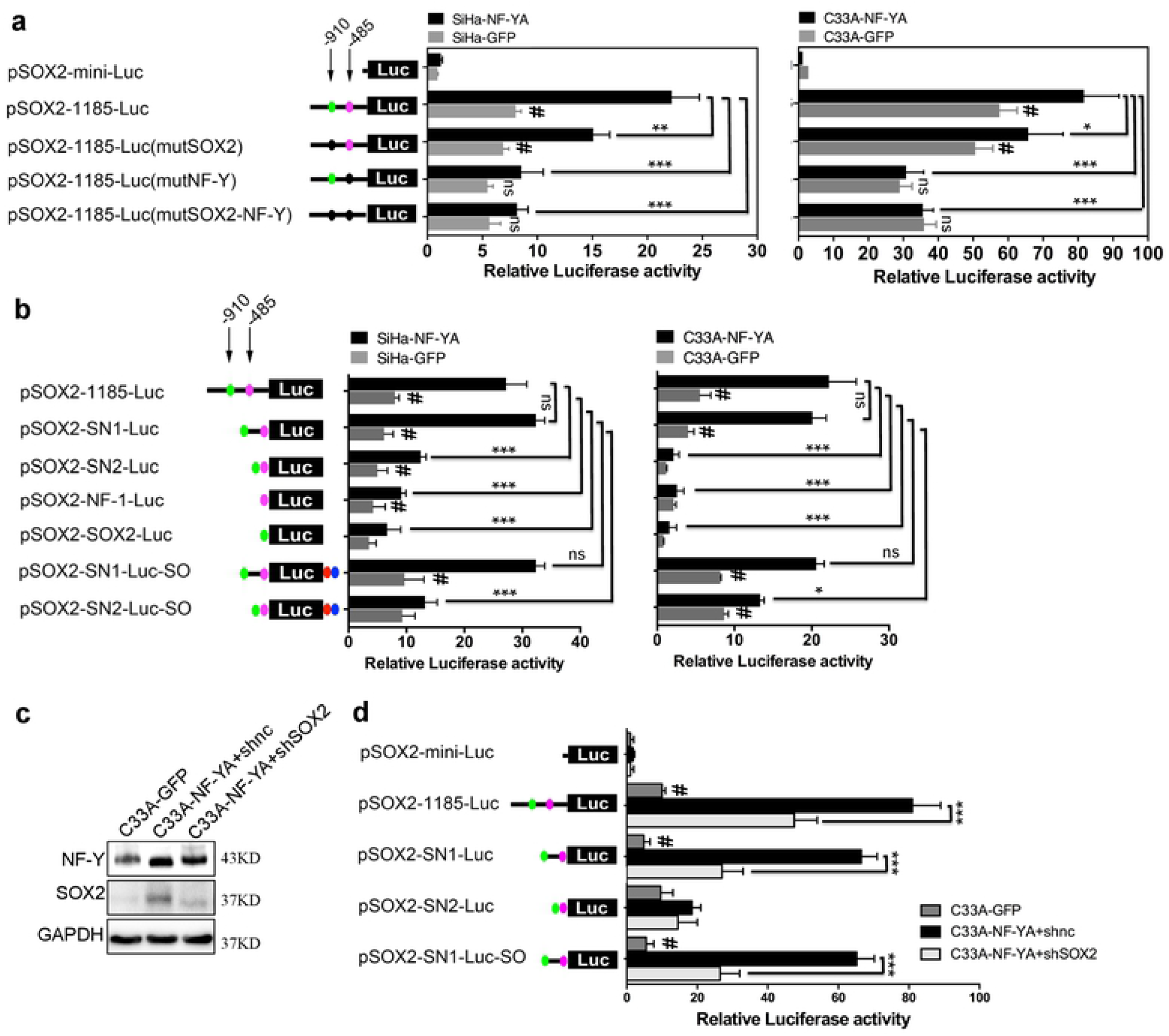
NF-Y specifically bound to CCAAT/ATTGG box upstream of the SOX2 promoter in cervical CSCs. (a) The luciferase activity of the SOX2 promoter mutations (green dot: SOX2 binding site; pink dot: NF-Y binding site; black dot: mutant of SOX2 or NF-Y binding site) in NF-YA-overexpressing SiHa and C333A cells. (b) The luciferase activity of the SOX2 promoter deletions in NF-YA-overexpressing SiHa and C333A cells. SOX2 was silenced by specific shRNA targeting the SOX2 CDS region in NF-YA-overexpressing C33A cells (c) and the luciferase activity of SOX2 promoter deletions was measured (d). (e) The diagram of NF-YA and SOX2 binding sites upstream of the SOX2 promoter. Data is presented as the mean ± SD of experiments in triplicate and statistically analyzed with student’s *t*-test. The symbols represent the following: *, *p* < 0.05; ***, *p* < 0.001; #, NF-YA group *vs* GFP group and *p* < 0.05; ns = no statistical difference.

## Discussion

Expression of transcription factor SOX2 is one of the hallmarks of embryonic stem cells, induced pluripotent stem cells and CSCs. The relationship between the expression of SOX2 and CSCs population was identified in breast cancer[12], lung cancer[15–17], ovarian cancer[18–20], and cervical cancer [21]. In our previous study, SOX2-positive cells isolated from the cervical cancer cell lines SiHa and C33A were found to exhibit self-renewal, differentiation, and tumor initiating properties, which are major characteristics of CSCs[7].

However, the key regions within the SOX2 promoter involved in the SOX2 transcription activity and how it maintains and regulates the CSC characteristics are not fully understood. Based on the on line promoter searcher system and known key region of SOX2 promoter in the murine and human embryo stem cell, we systematic analyzed the transcriptional elements, including the 11.5 kb of the human SOX2 promoter sequence. SOX2 promoter deletions and dual-luciferase reporter assay showed the crucial region of +1185 and −1600 was enough to activate the expression of SOX2 in the cervical CSC, a finding different from the 2 positive regulatory regions present within the SOX2 promoter region between −528 and +238 in ES cells[22]. Additionally, the presence of the downstream region −1600 seems too slightly enhanced but initiate the expression of SOX2. This suggests that the regulation of the endogenous SOX2 gene observed during the stemness maintenance of cervical CSCs is due to differential utilization of cis-regulatory elements in the region +1185 of SOX2 promoter.

To further define the cis-regulatory elements of this gene, the TFSEARCH database was used to examine the candidate sequence of the SOX2 promoter for possible transcription factor binding sites. This analysis identified a putative GAACAATG, CCAAT and ATGCATAT motif, which were the binding sites of SOX2, NF-Y and OCT4 factors, respectively. Mutagenesis of these motifs demonstrated that CCAAT box plays a crucial functional role in the regulation of the SOX2 promoter.

NF-Y (also known as CBF), a ubiquitously expressed trimetric transcription factor, has a dual role as both an activator and a repressor of transcription[23]. NF-Y regulates activity of target genes through a CCAAT box, a widespread control element mapping to proximal promoters, tissue-specific enhancers, and selected subclasses of human endogenous retrovirus (HERV) long terminal repeats (LTR). It is heterodimer protein complex that comprises three subunits (NF-YA, NF-YB and NF-YC). NF-YA is considered the limiting regulatory subunit of the trimer, since it is required for the complex assembly and sequence-specific DNA binding.

NF-Y has previously been identified as the marker of CSCs in hepatocellular carcinoma and embryonic carcinoma cells[24–26]. It also functions as the oncogene or suppressor in several carcinomas through a transcriptional mechanism in cell proliferation, metastasis and other malignant biological function[27, 28]. NF-Y was shown to regulate the expression of several human SOX genes, including SOX2, SOX3, SOX9, and SOX18[29–31]. This transcriptional activation function of NF-Y is mediated, at least in part, by direct binding to CCAAT boxes within promoters of target genes and by making complex interplay with other factors involved in transcriptional regulation of human SOX genes. Here, we found a new binding site for NF-Y in the SOX2 promoter, which was essential for the transcription of SOX2 protein in cervical CSCs.

In summary, we show that the NF-Y binding site CCAAT within the proximal region of the human SOX2 gene promoter plays a key role in regulating SOX2 expression in cervical CSCs. These results establish that NF-YA underlies SOX2 upregulation and is essential for the maintenance of characteristics of CSCs. We believe that these studies provide important insights into the biology of CSCs and identified NF-YA as a potential target for intervention in cervical cancer.

## Acknowledgements

This work was supported by a grant to Dr. Wen-Ting Yang from Basic Research Program of Natural Science in Shaanxi Province (2018JM7126114) and the National Natural Science Foundation of China (No. 81302278).

